# Reward-based improvements in motor sequence learning are differentially affected by dopamine

**DOI:** 10.1101/2023.03.02.530811

**Authors:** Sebastian Sporn, Joseph M Galea

**Author notes:** <u>Correspondence:</u> Sebastian Sporn.

## Abstract

Reward is a powerful tool to enhance human motor behaviour with research showing that it promotes motor sequence learning through increases in both motor vigour and movement fusion. Specifically, during a sequential reaching movement, monetary incentive leads to increased speed of each movement (vigour effect), whilst reward-based performance feedback increases speed of transition between movements (fusion effect). Therefore, motor sequence learning can be driven by distinct reward types with dissociable underlying processes. The neurotransmitter dopamine has been implicated to modulate motor vigour and regulate movement fusion. However, in humans, it is unclear if the same dopaminergic mechanism underlies both processes. To address this, we used a complex sequential reaching task in which rewards were based on movement times (MT). Crucially, MTs could be reduced via : 1) enhanced speed of individual movements (vigour effect) and/or 2) enhanced speed of transition between movements (fusion effect). 92 participants were randomly assigned to a reward and no reward group and were given either 2.5mg of the dopamine antagonist haloperidol or a placebo. Our results demonstrate that haloperidol impaired the reward-based effects on motor vigour whilst not affecting movement fusion. Thus, we illustrate that whilst both strategies are reward sensitive, they rely on dissociable mechanisms.

## Introduction

Ubiquitous in daily life, but often impaired in clinical populations^1–3^, the execution of motor sequences underlies a vast range of behaviours such as drinking a cup of coffee to driving a car. Within this context, reward has been demonstrated to promote motor sequence learning with research showing that enhanced motor sequence learning allows for a faster, yet similarly accurate execution of the motor sequence^4–9^. Therefore, reward may represent a powerful tool to promote motor sequence learning in rehabilitation settings. Importantly, recent work found that these reward-based improvements in motor sequence learning indexed as a shift in the speed-accuracy trade-off are driven by dissociable reward types^10,11^. Specifically, in a complex sequential reaching task, improvements in motor sequence learning (i.e., reduction in movement time – MT) were driven by two reward-based processes: 1) enhanced reaching speed of individual movements (represented by an increase in peak velocities) leading to a rapid decrease in MT which has also been observed in saccadic and simple discrete reaching tasks^12–18^ (vigour effect) and 2) a training-related increase in transition speed between movements reducing dwell times inside targets (movement fusion effect)^10^. Movement fusion represents a learning-dependent optimisation process during which individual motor elements of a sequence are blended into a combined singular action^10,19–21^. Therefore, movement fusion leads to decreases in MTs through a reduction in dwell times when transitioning between reaching movements. Importantly, fusion has not only been shown represent improved motor sequence learning indexed as increases in MTs but additionally has been demonstrated to lead to improvement in movement quality through enhanced smoothness/decreased jerk^10,19–21^. Thus, movement fusion represents a hallmark of sequential learning in which movements are combined in order to improve speed and efficiency^22^. Note here that movement fusion is conceptually close to the definitions of chunking^2,23–26^ and coarticulation^19–21,27–29^ with all three concepts describing improvements in movement speed and quality when transitioning between movement elements. Crucially, previous research has established that while a monetary incentive (providing money independent of performance feedback) enhanced motor vigour, reward-based performance feedback (providing feedback/points independent of money) improved movement fusion^10^. Therefore, reward may have motivational and informational properties that can both promote motor sequence learning. However, it is unclear if these two reward-based processes have dissociable neural mechanisms, which is of vital importance for reward to be employed in a strategic and targeted manner in clinical settings such as rehabilitation.

It has long been established that the neurotransmitter dopamine (DA) plays a central role in the processing of reward signals^30–32^ and has been suggested to modulate reaching speed (vigour)^33,34^, with experimental evidence coming from both animal^35^ and human^36–39^ work. Additionally, DA has been implicated to underlie movement fusion (chunking) with evidence coming from animals^40–43^, healthy humans^44^ and Parkinson’s patients^1,45,46^. Therefore, DA appears to play a crucial role in two core aspects of motor sequence learning which have also been shown to be reward sensitive. Importantly, while changes in tonic DA are believed to underlie the modulation of motor vigour, phasic DA has been suggested to underpin movement fusion (chunking)^34,47^. Within this context, it may seem plausible that the reward-based processes driven by monetary incentives and performance-based feedback are indeed based on dissociable dopaminergic mechanisms. However, to date this hypothesis has not been experimentally tested in humans.

To address this, we used a complex sequential reaching task in which participants were asked to execute a continuous sequence of eight reaching movements^10^. Motor vigour was measured as changes in peak velocities, while movement fusion was measured as changes in minimum velocities and alternatively as changes in the fusion index (FI) (see Methods for more information on FI). Participants received a combination of two rewards (i.e., monetary incentive and performance-based feedback) that have previously shown to distinctively enhance motor sequence learning by enhancing peak velocity of each movement (vigour effect) and reducing dwell times by increasing minimum velocities when transitioning between reaching movements (fusion effect). To investigate whether these reward-based processes rely on dissociable dopaminergic mechanisms, participants were randomly assigned to a reward and no reward group and were given either 2.5mg of the D2 antagonist haloperidol or a placebo. Our results replicated previous findings showing that both vigour and fusion were reward sensitive, with the reward groups showing significant increases in peak and minimum velocities and FI^10^. Importantly, our results demonstrate that haloperidol impaired the reward-based effects on motor vigour which was seen in participants showing lower peak velocities. In contrast, haloperidol did not affect movement fusion. Therefore, our results demonstrate that the reward-based processes driven by monetary incentives and performance-based feedback are indeed based on different mechanisms.

## Results

To investigate whether the reward-based improvement of both vigour and fusion relies on the same dopaminergic mechanism, we asked 92 participants to make 8 sequential reaching movements to designated targets (1 trial) using a motion tracking device (Figure 1a, b). Specifically, participants were randomly assigned to one of the four groups: Halo-Rew (N=25), Halo-NoRew (N=23), Placebo-Rew (N=24), Placebo-NoRew (N=23) (Figure 1c). Participants in the Halo groups received 2.5mg of the D2-antagonist haloperidol 120min prior to the start of the experiment coinciding with peak plasma concentration, while participants in the Placebo groups received a Placebo (lactose tablets). Importantly, while haloperidol is a known D2 antagonist it may also act as sedative which could influence results. Consequently, we included a self-report asking participants (N=42) to rate their perceived levels of fatigue and attention at the end of Day1 (drug intake). Analysis of the self-reported data did not reveal any significant group difference for both fatigue (Wilcoxon test: Z = 0.69, p = 0.4935, η^2^ = 0.11) and attention levels (Z = -1.05, p = 0.2955, η^2^ = 0.16). These results highlight that any changes in performance due to haloperidol can most likely be attributed to it modulating dopamine levels. Furthermore, participants in the Rew groups were able to earn money during training depending on how fast they complete a trial and received feedback on their earnings after each trial (Figure 1d left). Therefore, Rew participants received a combination of a monetary incentive and performance-based feedback. Importantly, participants could pursue two strategies to reduce MTs in this task: 1) increases in peak velocities of the individual movements (vigour effect) and 2) reductions in dwell-times when transitioning between movements via increases in minimum velocities (fusion effect; note here that these strategies are not mutually exclusive and can be combined to further reduce MTs - Figure 1e). NoRew participants were instructed to complete each trial as fast and accurately as possible (Figure 1d right). Prior to the start of the experiment, participants were trained on the sequence without a time constraint until reaching a learning criterion of 5 successful trials in a row. Importantly, missing a target resulted in an immediate abortion of the current trial, which participants then had to repeat. Participants then completed a baseline period (10 trials) during which they were encouraged to complete each trial ‘as fast and as accurately as possible’ (Figure 1f). Afterwards, participants in each group completed 200 training trials before engaging in a rewarded (post-R) and no rewarded (post-NR) post-assessment (20 trials each). Therefore, all groups received both monetary incentive and performance-based feedback during post-R, while neither was available during post-NR.

**Figure 1.**
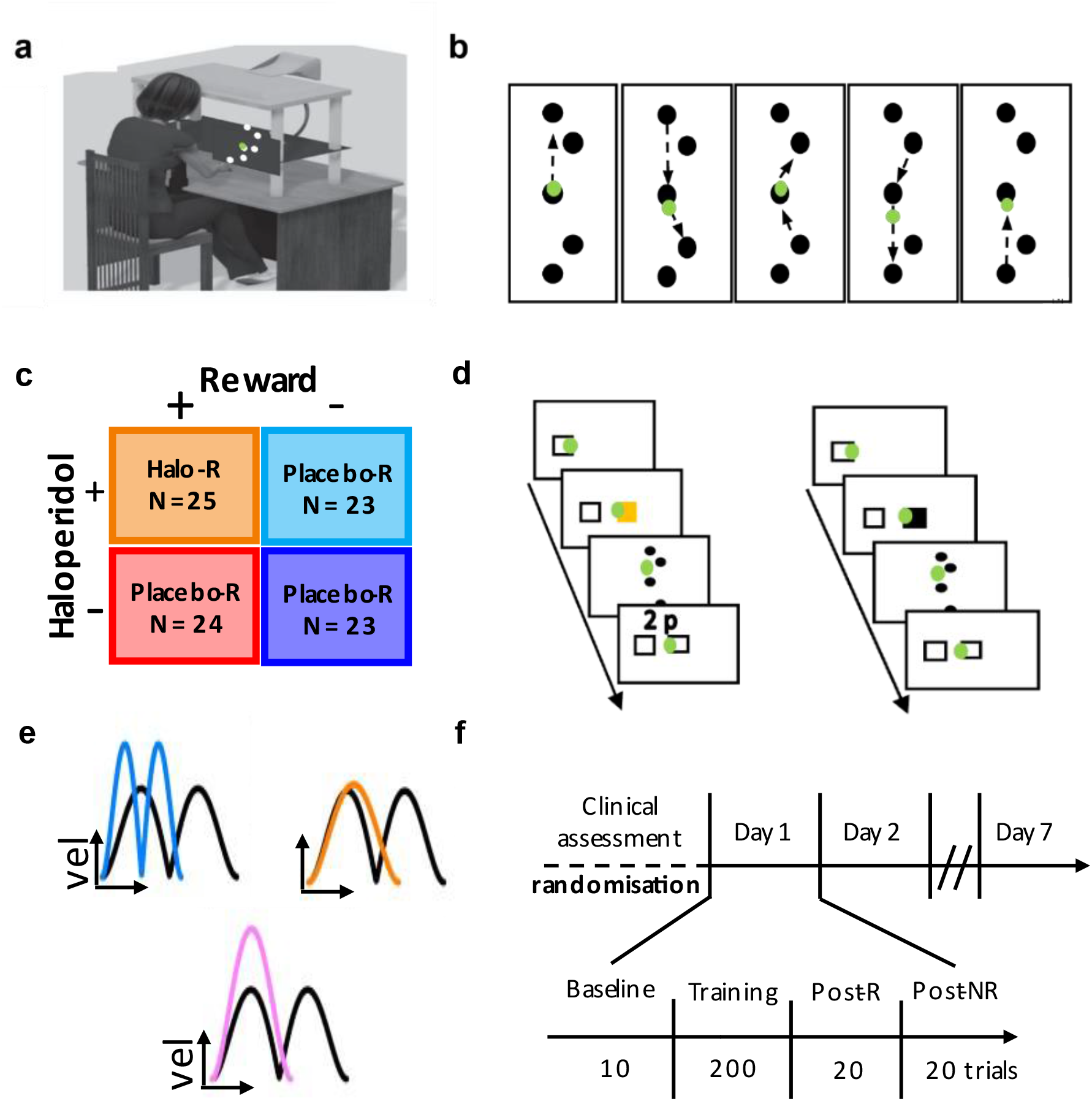
Experimental setup. **A)** Participants wore a motion-tracking device on the index finger and the unseen reaching movements were performed across a table whilst a green cursor matching the position of index finger was viewable on a screen. **B)** 8 movement sequential reaching task. Participants started from the centre target. **c)** Groups. Participants either received monetary incentive with accurate performance-based feedback (R) or not (NR). Similarly, participants were either received 2.5mg of haloperidol (Halo) or a placebo (Placebo). **d)** Trial design. Reward trials (left) were cued using a visual stimulus (yellow start box) prior to the start of the trial and participant received performance-based feedback depending on how fast they completed the trial. In contrast, no reward trials were not cued, and no feedback was given (right). Instead, participants were asked to complete each trial as fast and accurately as possible. **e)** Strategies. To reduce MTs to earn more money participants could pursue two independent strategies: 1) Increase peak velocities (top right) and 2) Fuse consecutive movements leading to increases in minimum velocities (top left). Note these strategies are not mutually exclusive and can be combined to further reduce MTs (bottom). **f)** Study design. Prior to the start of the experiment, participants were trained on the reaching sequence and were then asked to perform 10 baseline trials. Randomly allocated to one of four groups, participants completed 200 trials during training and an additional 20 trials in each post assessment; one with reward (post-R) and one without (post-NR). This design was repeated on Day2 (without drug intake) and on Day7 participants engaged in a post-R and post-NR (25 trials).

The post-assessment intended to compare performance between groups when under the same condition (Figure 1f). The same experimental design was repeated on Day2 without any drug/placebo intake, to ascertain that potential haloperidol effects found on Day1 are indeed attributable to a drug manipulation and is therefore not present on Day2. Additionally, participants returned on Day7 to complete a rewarded (post-R) and no rewarded (post-NR) post-assessment (25 trials each).

### Haloperidol modulates motor sequence learning only on Day1

Enhanced motor sequence learning was indexed as a reduction in MTs with MT reflecting total movement duration from exiting the start box until reaching the last target (Figure 2a). Using a Kruskal Wallis test, with group as a factor, we did not find any significant differences at Baseline (F = 5.44, p = 0.1424). We then conducted a linear mixed-effect model (LMM) to assess the effect of drug, reward and trial number on MT performance over the course of training. To this end, we included MT as the dependent variable and reward, drug and trial number as fixed effects (see Methods for more information on the linear mixed models). We found a significant main effect for trial number (Table 1) suggesting that over Day1 MTs decreased over the course of training. Similarly, reward was significant, which replicated previous results showing that a monetary incentive in combination with accurate performance-based feedback promotes motor sequence learning as seen in a greater reduction in MTs^10^. Importantly, we also found a significant interaction between trial number and drug (Table 1).

**Table 1.** Mixed-effect model for MT for Day1.

| Variable | Estimate | SE | df | t | p |
| --- | --- | --- | --- | --- | --- |
| Intercept | 4.464 | 0.098 | 88 | 45.353 | < .001 |
| Trial | -0.275 | 0.047 | 88 | -5.820 | < .001 |
| Reward | -0.654 | 0.098 | 88 | -6.648 | < .001 |
| Drug | -0.119 | 0.098 | 88 | -1.206 | 0.231 |
| Trial * Reward | 0.072 | 0.047 | 88 | 1.517 | 0.133 |
| Trial * Drug | 0.106 | 0.047 | 88 | 2.251 | **0.027** |
| Reward * Drug | -0.033 | 0.098 | 88 | -0.339 | 0.736 |
| Trial * Reward * Drug | 0.076 | 0.047 | 88 | 1.612 | 0.110 |

**Figure 2.**
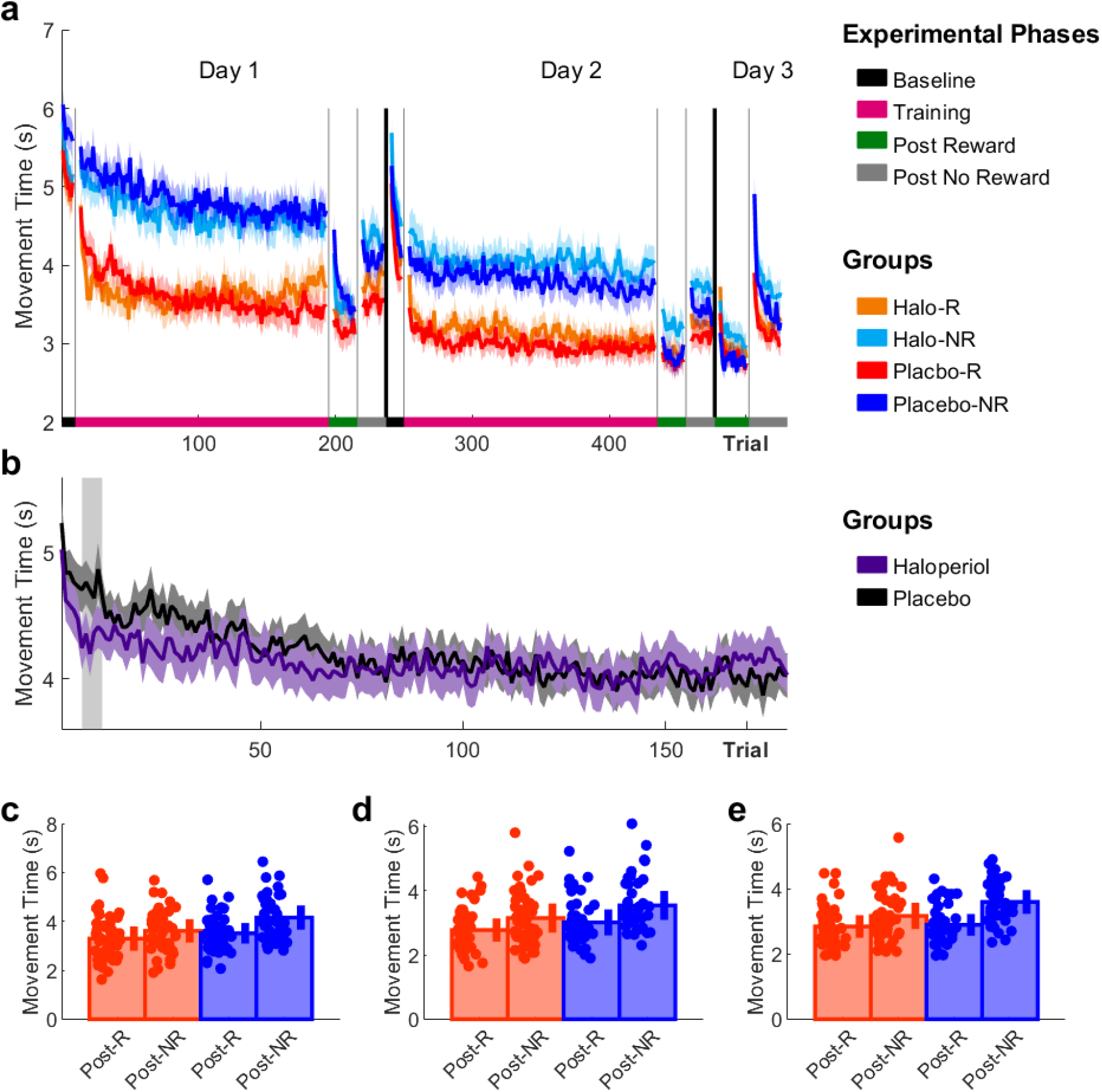
Haloperidol leads to a global slowing in MTs only on Day1. **A)** Trial-by-trial changes in MT averaged over participants for all groups. **b)** Trial-by-trial changes in MT averaged over both Halo and Placebo groups. Regions shaded in grey indicate significant differences in performance between groups **c-e)** Post assessment performance (post-R vs post-NR) comparing Rew and NoRew groups on **c)** Day1, **d)** Day2 and **e)** Day7. Shaded regions/error bars represent SEM.

To further investigate this effect of drug over time, we collapsed the Halo and Placebo groups and ran a permutation analysis to investigate at which time point these two groups significantly differ. For the permutation test, statistical significance was established using a row-shuffle non-parametric permutation scheme. To this end, 1000 permutations were computed for each time point across all participants before statistical significance was determined at p < 0.05 (see Methods for more information on the permutation analysis. Interestingly, we found that the Halo groups were significantly faster during early training (Figure 2b). However, this effect was limited to Trial 6-11 suggesting that over the course of training they progressively slowed down and showed higher MTs when compared to the Placebo groups towards the end of training. Note here that this difference in MT performance was not significant towards the end of training, yet it suggests that haloperidol led to a slowing which was not apparent in the Placebo groups (Figure 2b).

Importantly, a separate LMM conducted for Day2 did not reveal a significant Trial x Drug interaction which highlights that the slowing observed in the Halo groups on Day1 is indeed driven by the DA manipulation (Table 2). Instead, we again found a significant main effect for both Trial and Reward. These results suggest that MT performance further improved over the course of training on Day2 and that the Rew groups exhibited faster MTs when compared to the NoRew groups.

**Table 2.** Mixed-effect model for MT for Day2.

| Variable | Estimate | SE | df | t | p |
| --- | --- | --- | --- | --- | --- |
| Intercept | 3.624 | 0.085 | 88.022 | 42.813 | < .001 |
| Trial | -0.141 | 0.032 | 88.019 | -4.394 | < .001 |
| Reward | -0.437 | 0.085 | 88.022 | -5.158 | < .001 |
| Drug | 0.087 | 0.085 | 88.022 | 1.024 | 0.309 |
| Trial * Reward | 0.007 | 0.032 | 88.019 | 0.231 | 0.818 |
| Trial * Drug | 0.005 | 0.032 | 88.019 | 0.161 | 0.873 |
| Reward * Drug | 0.002 | 0.085 | 88.022 | 0.021 | 0.983 |
| Trial * Reward * Drug | -0.023 | 0.032 | 88.019 | -0.704 | 0.483 |

Assessing performance across post assessment, we conducted separate mixed-effect ANOVAs for each day (i.e., Day1, Day2 and Day7) with reward (R vs NR) and drug (Halo vs Placebo) as between factors and timepoint (post-R vs post-NR) as a within factor. While the effect size of significant findings were represented by η^2^, significant interactions were further analysed using Wilcoxon tests with false discovery rate (FDR) being used to correct for multiple comparisons. For Day1 and Day2, we found a significant main effect for reward (Day1: F = 5.15, p = 0.0257, η^2^ = 0.84; Day2: F = 4.80, p = 0.0310, η^2^ = 0.83) with Day3 being close to significance (Day3: F = 3.71, p = 0.0573, η^2^ = 0.79). Furthermore, timepoint was significant for all days (Day1: F = 68.48, p < 0.0001, η^2^ = 0.99; Day2: F = 65.07, p < 0.0001, η^2^ = 0.99; Day3: F = 100.22, p < 0.0001, η^2^ = 0.99). Importantly, we also found a significant interaction between reward and timepoint for Day1 and Day 3 (Day1: F = 8.20, p = 0.0052, η^2^ = 0.89, Figure 2c; Day2: F = 1.93, p = 0.1681, η^2^ = 0.66, Figure 2d; Day3: F = 13.54, p < 0.0001, η^2^ = 0.93, Figure 2e).

Post-hoc analysis for Day1 revealed a significant change in MT performance from post-R to post-NR only for the NoRew groups (Z = -3.25, p = 0.0047, η^2^ = 0.35) but not the Rew groups (Z = -1.90, p = 0.0622, η^2^ = 0.20, Figure 2c). This suggests that MT performance became more stable and less reward dependent in the Rew groups. This is further supported by the significant MT difference between the groups when comparing performance in post-NR (Z = -2.67, p = 0.0152, η^2^ = 0.28). Additionally, the three-way interaction between reward x drug x timepoint almost reached significance (F = 3.7, p = 0.0570, η^2^ = 0.79) indicating that the drug may also act on much smaller time scales with changes in the reward structure after only 20 trials. Post hoc analysis for Day3 revealed a significant increase in MTs from post-R to post-NR for the NoRew groups (Z = -4.60, p < 0.0001, η^2^ = 0.50) and also Rew groups (Z = -2.23, p = 0.0342, η^2^ = 0.23). Additionally, we found a significant difference in MT between groups for post-NR (Z = -3.10, p = 0.0038, η^2^ = 0.32) but not post-R (Z = - 0.56, p = 0.5758, η^2^ = 0.06, Figure 2d). This suggests that performance in the Rew groups was less reward-dependent when compared to the NoRew groups, but nevertheless was less stable than on Day1 and 2.

### Haloperidol impairs reward-based effects on movement vigour

In this task, participants could decrease MTs through two independent strategies: 1) increases in peak velocities (vel_peak_, vigour effect) and 2) reductions in dwell-times between reaches via increases in minimum velocities (vel_min_, fusion effect) (Figure 1e). To ascertain whether these strategies are indeed independent, we first conducted a stepwise linear regression with MT as the dependent variable and vel_peak_, vel_min_ and movement fusion measured as a fusion index (FI) as covariates. The full model accounted for 84.6% of the variance found (adjusted R^2^ = 0.846, see Supplementary Table 1). However, the model greatly violated the multicollinearity criterion, with an average variance inflation factor (VIF) of 12.022 (see Supplementary Table 2). This could be expected because FIs proportionally increase with increasing vel_mins_ when vel_max_ are held constant. However, the model with just vel_peak_ and vel_min_ as covariates accounted for 73.7% of the variance (adjusted R^2^ = 0.737) and did not violate the multicollinearity criterion (average VIF of 1.92 and a tolerance of 0.521). Importantly, both factors were highly significant (vel_max_: t = -132.01, p < 0.0001; vel_min_: t = -107.01, p < 0.0001, see Supplementary Table 2). These results highlight that both strategies reduce MTs whilst not being highly correlated with each other.

Vel_peak_ represents the average of maximum velocities achieved during the eight reaching movements to complete a trial. Yet again, we found no differences at Baseline (ANOVA; group: F = 5.79, p = 0.1223). We then conducted a separate LMM for Day1 and Day2 with vel_peak_ as the dependent variable and reward, trial and drug as covariates. Similarly to MT, we found a significant main effect for both reward and trial on Day1 and a significant interaction between them (Table 3; Figure 3a). These results yet again replicate previous findings highlighting that reward enhances movement vigour^10,14^.

**Table 3.**
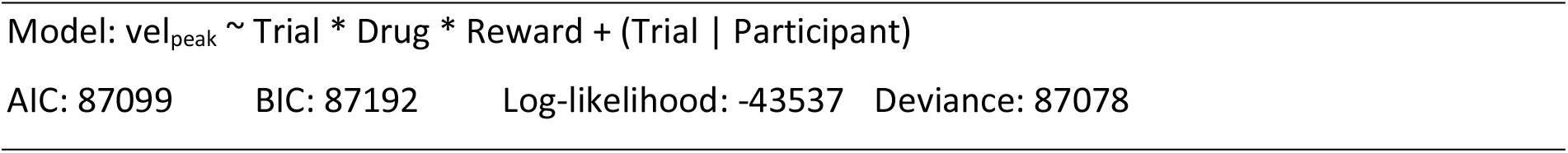
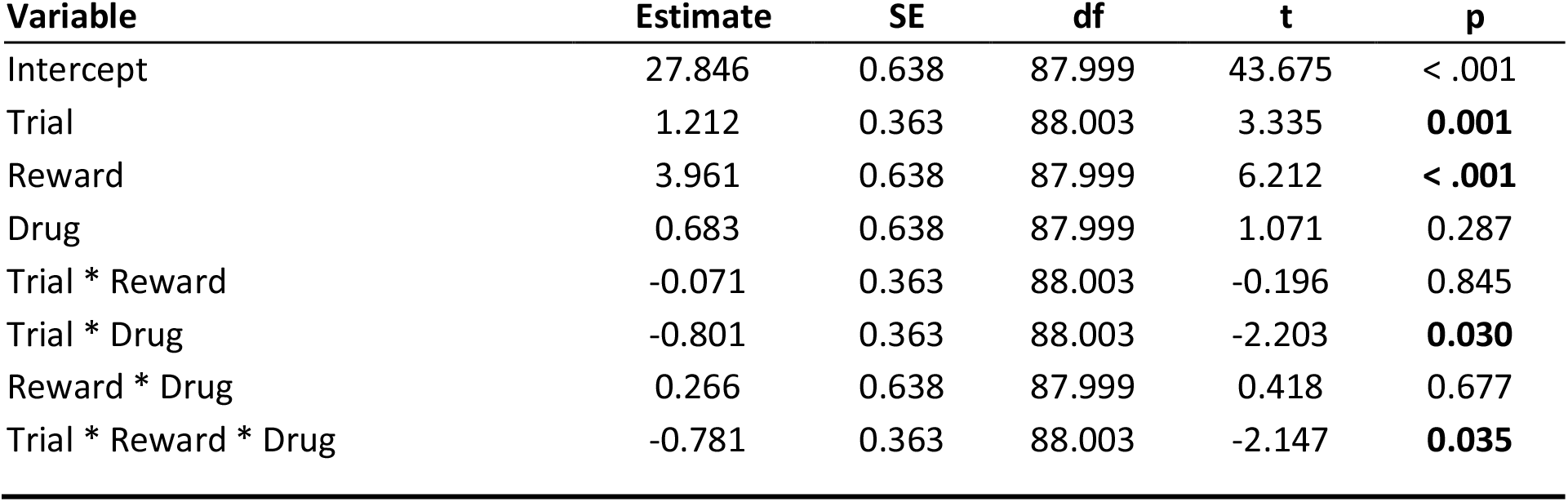
Mixed-effect model for vel_peak_ for Day1.

| Variable | Estimate | SE | df | t | p |
| --- | --- | --- | --- | --- | --- |
| Intercept | 27.846 | 0.638 | 87.999 | 43.675 | < .001 |
| Trial | 1.212 | 0.363 | 88.003 | 3.335 | **0.001** |
| Reward | 3.961 | 0.638 | 87.999 | 6.212 | **< .001** |
| Drug | 0.683 | 0.638 | 87.999 | 1.071 | 0.287 |
| Trial * Reward | -0.071 | 0.363 | 88.003 | -0.196 | 0.845 |
| Trial * Drug | -0.801 | 0.363 | 88.003 | -2.203 | **0.030** |
| Reward * Drug | 0.266 | 0.638 | 87.999 | 0.418 | 0.677 |
| Trial * Reward * Drug | -0.781 | 0.363 | 88.003 | -2.147 | **0.035** |

**Figure 3.**
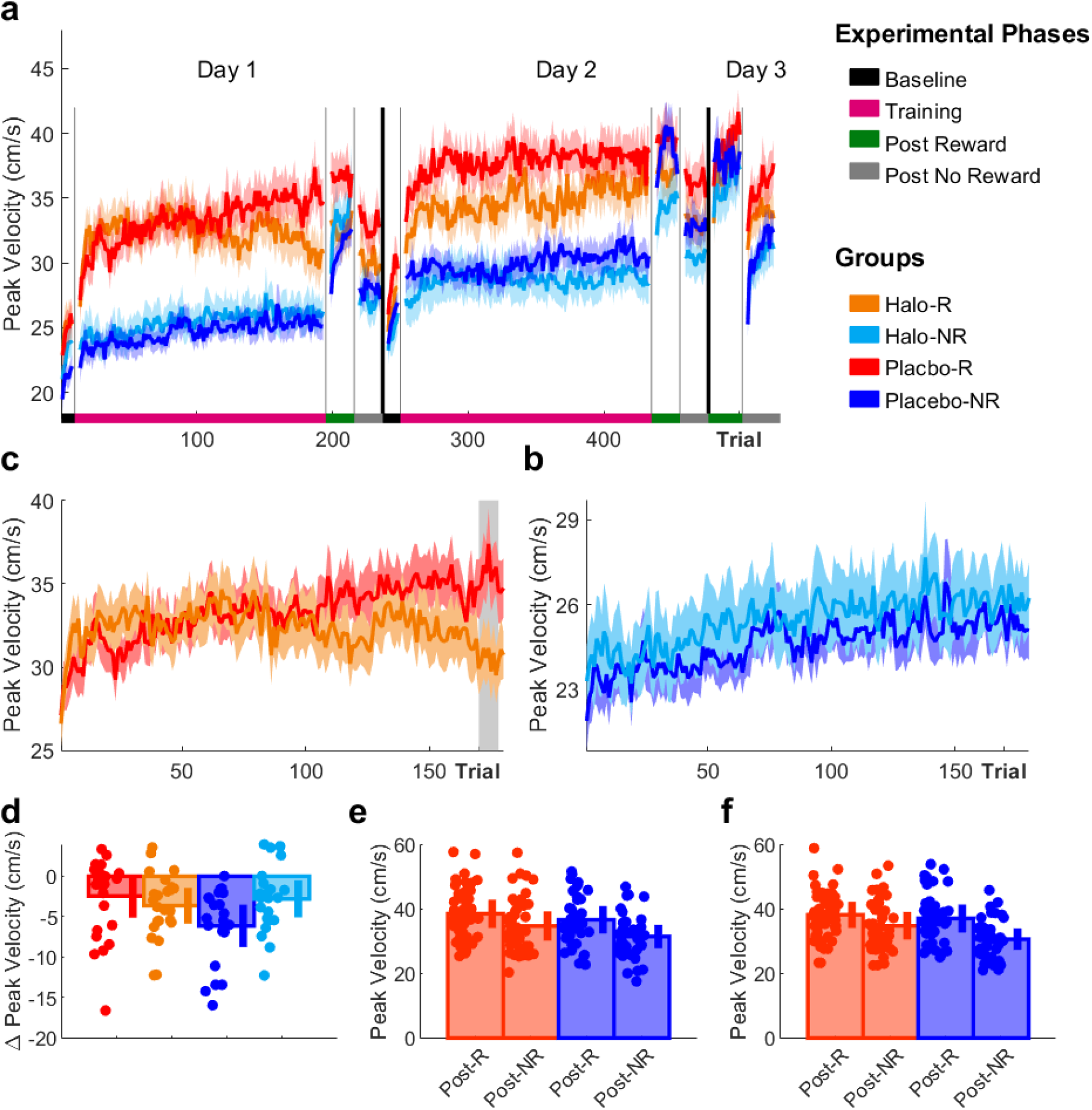
Haloperidol leads to a selective decrease in peak velocities only for the Halo-R group on Day1. **A)** Trial-by-trial changes in vel_peak_ averaged over participants for all groups. **b-c)** Trial-by-trial changes in vel_peak_ for **b)** Halo-R and Placebo-R and **c)** Halo-NR and Placebo-NR for Day1. Regions shaded in grey indicate significant differences in performance between groups **d-f)** Post assessment performance (post-R vs post-NR) comparing **d)** differences in vel_peak_ from post-R to post-NR between all groups on Day1 **e)** differences across all groups between timepoints (left) and across timepoints between Halo and Placebo groups (right) on Day2, and **e)** differences in vel_peak_ between Rew and NoRew groups on Day7. Shaded regions/error bars represent SEM.

Importantly, we found a significant three-way interaction between reward, trial, and drug. We further investigated this interaction by running separate permutation analyses for the reward (Figure 3b) and no reward groups (Figure 3c) to investigate at which time point the halo and placebo groups significantly differed. We found that the Halo-R group exhibited significantly lower vel_peak_ towards the end of training on Day1 when compared to Placebo-R (Trial 170-180), even though from visual inspection they had higher vel_peak_ in the beginning of training (Figure 3b). Crucially, we did not observe any performance differences between the NoRew groups (Figure 3c), which indicates that haloperidol selectively impaired the reward-based effects on vigour. Converging results come from the LMM conducted separately for Day2, where we did not find a significant three-way interaction between reward, trial, and drug (Table 4). Instead, we found significant main effects for both reward and trial on Day2.

**Table 4.**
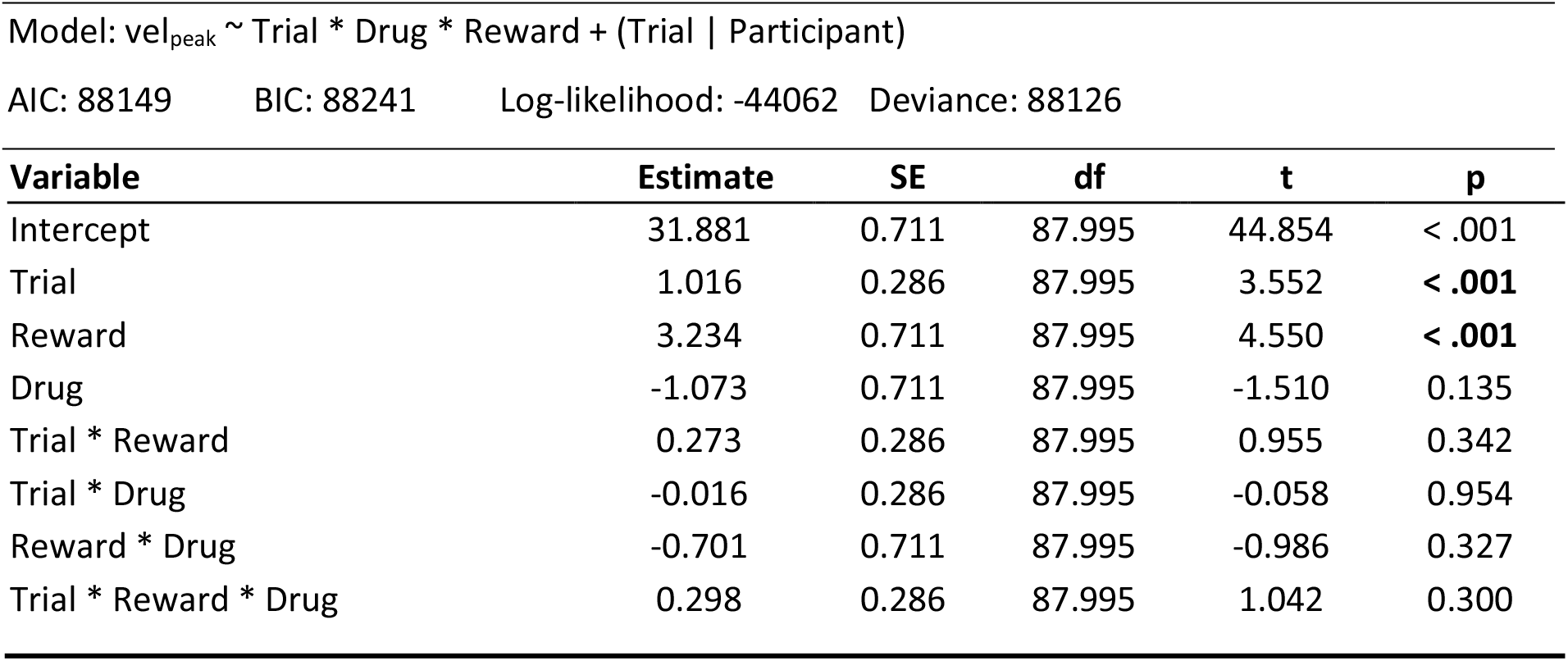
Mixed-effect model for vel_peak_ for Day2.

Assessing performance across post assessment, we conducted separate mixed-effect ANOVAs for each day and found a main effect for reward (F = 5.01, p = 0.0277, η^2^ = 0.83) and timepoint (F = 65.94, p < 0.0001, η^2^ = 0.99) for Day1. Importantly, we also found a significant interaction between reward, timepoint and drug on Day1 (F = 5.97, p = 0.0166, η^2^ = 0.86). To further assess this result, we calculated the difference in average performance between post-R and post-NR for each subject and ran separate Wilcoxon tests for each pairing (N=4) and used FDR to correct for multiple comparisons (Figure 3d). This post-hoc analysis revealed a significant difference in vel_peak_ change from post-R to post-NR between Halo-R and Placebo-R (Z = 2.87, p = 0.0166, η^2^ = 0.42, Figure 3d). However, no further post-hoc test reached significance. Interestingly, on Day2 we found a significant main effect for drug (F = 4.27, p = 0.0418, η^2^ = 0.81, Figure 3e right) and yet again timepoint (F = 75.34, p < 0.0001, η^2^ = 0.99, Figure 3e left). For Day3, we yet again found a significant timepoint effect (Day3: F = 87.55, p < 0.0001, η^2^ = 0.99) and a significant reward x timepoint interaction (Day3: F = 8.17, p = 0.0053, η^2^ = 0.89, Figure 2f). Similarly to Day1, we found a significant increase in vel_peak_ from post-R to post-NR for the NoRew groups (Z = 3.80, p < 0.0001, η^2^ = 0.41) and also Rew groups (Z = 2.42, p = 0.0209, η^2^ = 0.24). Additionally, we found a significant difference in vel_peak_ between groups for post-NR (Z = 2.244, p = 0.0209, η^2^ = 0.25) but not post-R (Z = 0.96, p = 0.3377, η^2^ = 0.10, Figure 2f). This suggests that performance in the Rew groups was less reward-dependent when compared to the NoRew groups, but nevertheless showed an increase in vel_peak_ when moving to post-NR.

### Haloperidol does not impair movement fusion

In this task improved motor sequence learning indexed as a decrease in MTs could be driven by increases in vigour and fusion. To investigate whether haloperidol affects the reward-based effect on fusion, we assessed changes in vel_min_ and FI. Vel_min_ represents the average of minimum velocities achieved when transitioning between sequential reaching movements. They, therefore, represent a proxy measure of how long participants dwelled in via targets before executing a further reaching movement to arrive at the subsequent target (see Methods for further information). Yet again, we did not find any differences between groups at Baseline (ANOVA; Group: F = 3.68, p = 0.2977). We then conducted a separate LMM for Day1 and Day2 with vel_min_ as the dependent variable and reward, trial and drug as covariates. Similarly to the previous results, we found a significant main effect for both reward and trial on Day1 (Table 5; Figure 4a). These results yet again replicate previous findings highlighting that reward increases minimum velocities and leads to shorter dwell times in the via points^12^. Importantly, in contrast to vel_peak_, we did not find a significant drug effect or interaction with other predictors, which suggests that haloperidol does not impair this specific measure of movement fusion. Note here that previous research has shown that increases in vel_min_ are learning-dependent and lead to the expression of movement fusion, which will be looked at in the next section^10^.

**Table 5.** Mixed-effect model for vel_min_ for Day1.

| Variable | Estimate | SE | df | t | p |
| --- | --- | --- | --- | --- | --- |
| Intercept | 4.406 | 0.372 | 88.000 | 11.839 | < .001 |
| Trial | 1.237 | 0.235 | 88.005 | 5.276 | < .001 |
| Reward | 1.656 | 0.372 | 88.000 | 4.449 | < .001 |
| Drug | 0.575 | 0.372 | 88.000 | 1.545 | 0.126 |
| Trial * Reward | 0.373 | 0.235 | 88.005 | 1.590 | 0.115 |
| Trial * Drug | -0.370 | 0.235 | 88.005 | -1.578 | 0.118 |
| Reward * Drug | 0.396 | 0.372 | 88.000 | 1.064 | 0.290 |
| Trial * Reward * Drug | -0.217 | 0.235 | 88.005 | -0.926 | 0.357 |

**Figure 4.**
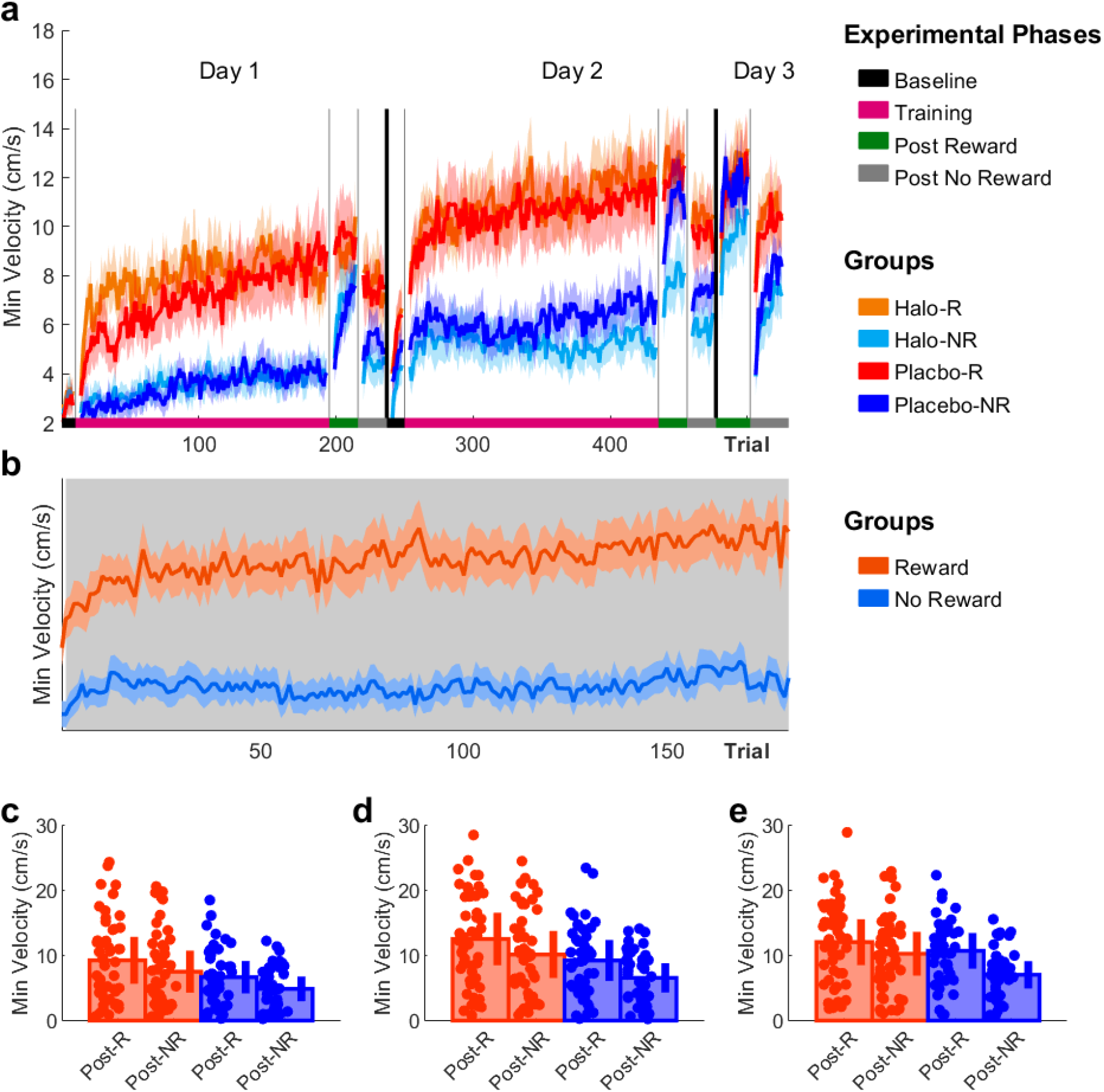
Haloperidol does not lead to an increase in minimum velocities. **A)** Trial-by-trial changes in vel_min_ averaged over participants for all groups. **b)** Trial-by-trial changes in vel_min_ across Rew and NoRew groups during Training on Day2. Regions shaded in grey indicate significant differences in performance between groups **c-e)** Post assessment performance (post-R vs post-NR) comparing Rew and NoRew groups on **c)** Day1, **d)** Day2 and **e)** Day7. Shaded regions/error bars represent SEM.

LMM results for Day2 revealed a significant main effect for reward and trial and also a significant interaction between these two (Table 6). To further investigate this effect of reward over time, we collapsed the Rew and NoRew groups and ran a permutation analysis to investigate at which time point these two groups significantly differ. We found that the Rew groups exhibited higher vel_min_ across the entire training on Day2 (only trials not significant at p < 0.05 were Trial 1 and 2, Figure 4b). Importantly, the result also suggests that the reward groups may further improve vel_min_ performance over the course of training which aligns with previous results showing that learning to transition between sequential reaching movements faster is a training-depending process that is reward sensitive.

**Table 6.** Mixed-effect model for vel_min_ for Day2.

| Variable | Estimate | SE | df | t | p |
| --- | --- | --- | --- | --- | --- |
| Intercept | 7.585 | 0.548 | 88.000 | 13.829 | < .001 |
| Trial | 0.723 | 0.200 | 87.998 | 3.618 | < .001 |
| Reward | 2.109 | 0.548 | 88.000 | 3.845 | < .001 |
| Drug | -0.043 | 0.548 | 88.000 | -0.078 | 0.938 |
| Trial * Reward | 0.461 | 0.200 | 87.998 | 2.307 | **0.023** |
| Trial * Drug | -0.086 | 0.200 | 87.998 | -0.430 | 0.668 |
| Reward * Drug | 0.151 | 0.548 | 88.000 | 0.276 | 0.783 |
| Trial * Reward * Drug | 0.219 | 0.200 | 87.998 | 1.094 | 0.277 |

Assessing performance across post assessment, we conducted separate mixed-effect ANOVAs for each day and found for each day a main effect for reward (Day1: F = 5.99, p = 0.0164, η^2^ = 0.87; Day2: F = 8.11, p = 0.0055, η^2^ = 0.89; Day7: F = 4.33, p = 0.0381, η^2^ = 0.82) and timepoint (Day1: F = 37.25, p < 0.0001, η^2^ = 0.97; Day2: F = 45.14, p < 0.0001, η^2^ = 0.98; Day7: F = 64.91, p < 0.0001, η^2^ = 0.99). Importantly, we also found a significant reward x timepoint interaction only for Day7 (Day1: F = 0.07, p = 0.8939, η^2^ = 0.02, Figure 4c; Day2: F = 0.90, p = 0.7082, η^2^ = 0.12, Figure 4d; Day7: F = 7.67, p = 0.0069, η^2^ = 0.87, Figure 4e). Post-hoc analysis revealed that vel_min_ changed significantly from post-R to post-NR only for the NoRew groups (Z = 3.50, p = 0.0019, η^2^ = 0.38, Figure 4e) but not for the Rew groups (Z = 1.51, p = 0.1736, η^2^ = 0.15). Additionally, performance was significantly different when comparing the Rew and NoRew groups during post-NR (Z = 2.64, p = 0.0165, η^2^ = 0.28) but not post-R (Z = 1.15, p = 0.2516, η^2^ = 0.12). These results suggest that vel_min_ performance was less reward dependent and became more stable in the Rew groups.

### Haloperidol does not impair motor sequence learning measured through fusion

Movement fusion describes the slow process of fusing discrete movements into single, continuous action^10,19–21^ which conceptually aligns with the concepts of chunking and coarticulation. It is characterised by increases in vel_min_ around the via point between reaching movements which leads to a reduction in dwell time. Therefore, movement fusion leads to decreases in MTs and crucially also to an improvement in movement quality considering that it is associated with enhanced smoothness/reduced jerk^10,19–21^. Thus, movement fusion represents a hallmark of sequential learning which has previously been shown to be reward sensitive. Here we measured movement fusion using FI with higher values for FI indicating more pronounced fusion of movements (see Methods for more information). Baseline analysis did not reveal any differences between groups in FI scores (ANOVA, Group: F = 2.66, p = 0.4478).

Similarly to the vel_min_ results, we found a significant main effect for both reward and trial on Day1 (Table 7; Figure 5a). These results yet again replicate previous findings highlighting that reward increases movement fusion and leads to shorter dwell times around the via points thereby decreasing MTs^12^. Importantly, in contrast to vel_peak_ we did not find a significant drug effect or interaction with other predictors, which suggests that haloperidol does not impair fusion. Similarly to the vel_min_ results, LMM results for Day2 revealed a significant main effect for reward and trial and also a significant interaction between these two (Table 8). To further investigate this effect of reward over time, we collapsed the Rew and NoRew groups and ran a permutation analysis to investigate at which time point these two groups significantly differ. We found that the Rew groups exhibited higher FI values across the entire training on Day2 (only trials not significant at p < 0.05 were Trial 1 and 2, Figure 5b).

**Table 7.** Mixed-effect model for FI for Day1.

| Variable | Estimate | SE | df | t | p |
| --- | --- | --- | --- | --- | --- |
| Intercept | 1.078 | 0.075 | 88.001 | 14.443 | < .001 |
| Trial | 0.227 | 0.038 | 88.003 | 6.039 | < .001 |
| Reward | 0.230 | 0.075 | 88.001 | 3.080 | **0.003** |
| Drug | 0.112 | 0.075 | 88.001 | 1.497 | 0.138 |
| Trial * Reward | 0.035 | 0.038 | 88.003 | 0.924 | 0.358 |
| Trial * Drug | -0.061 | 0.038 | 88.003 | -1.614 | 0.110 |
| Reward * Drug | 0.050 | 0.075 | 88.001 | 0.669 | 0.505 |
| Trial * Reward * Drug | 0.001 | 0.038 | 88.003 | 0.027 | 0.979 |

**Table 8.** Mixed-effect model for FI for Day2.

| Variable | Estimate | SE | df | t | p |
| --- | --- | --- | --- | --- | --- |
| Intercept | 1.626 | 0.094 | 88.000 | 17.363 | < .001 |
| Trial | 0.092 | 0.025 | 88.000 | 3.659 | < .001 |
| Reward | 0.266 | 0.094 | 88.000 | 2.838 | **0.006** |
| Drug | 0.028 | 0.094 | 88.000 | 0.300 | 0.765 |
| Trial * Reward | 0.051 | 0.025 | 88.000 | 2.017 | **0.047** |
| Trial * Drug | -0.005 | 0.025 | 88.000 | -0.211 | 0.833 |
| Reward * Drug | 0.076 | 0.094 | 88.000 | 0.815 | 0.417 |
| Trial * Reward * Drug | 0.028 | 0.025 | 88.000 | 1.107 | 0.271 |

**Figure 4.**
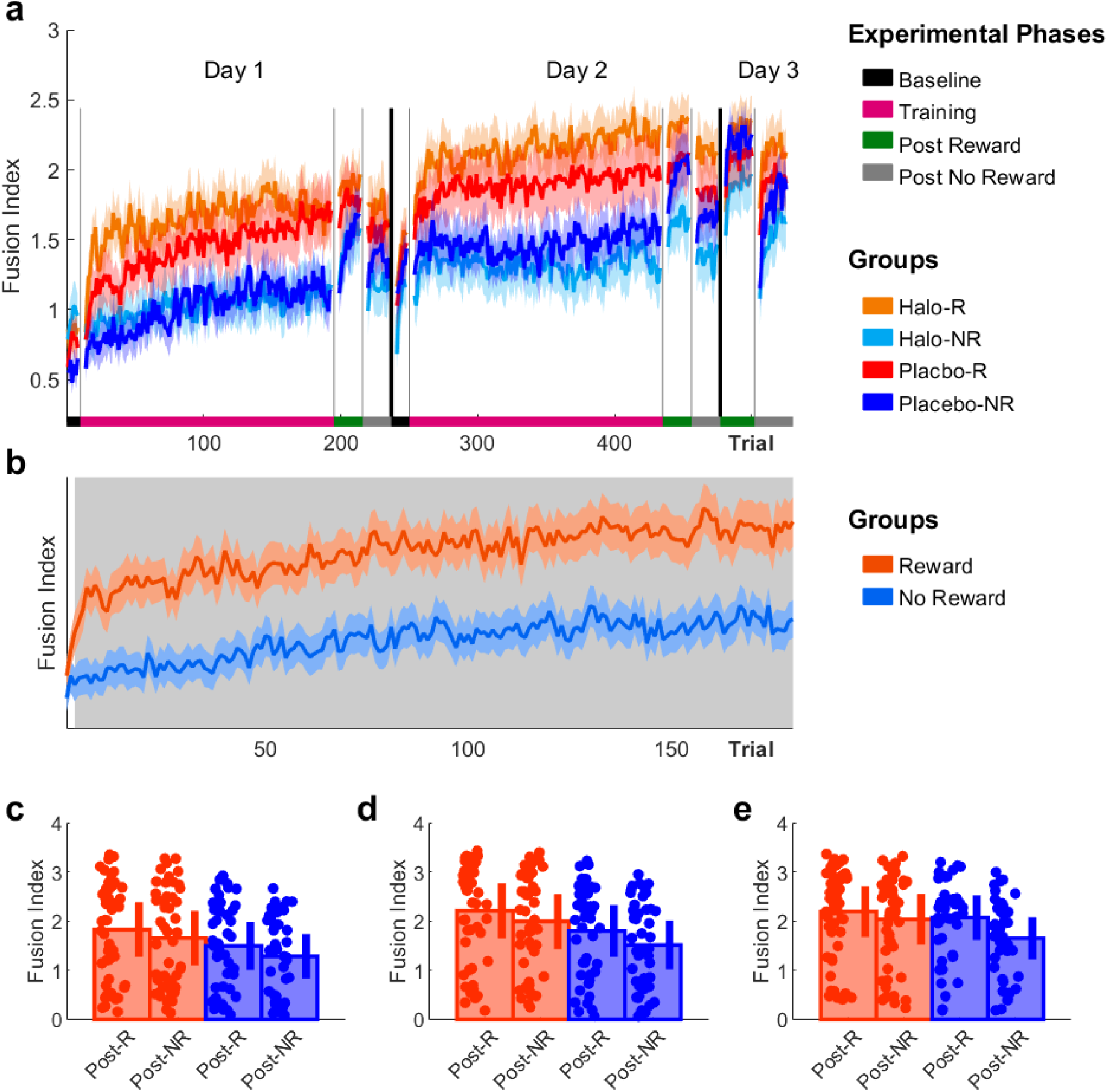
Haloperidol does not impair motor sequence skill learning measured in movement fusion. **a)** Trial-by-trial changes in FI scores averaged over participants for all groups. **b)** Trial-by-trial changes in FI scores across Rew and NoRew groups during Training on Day2. Regions shaded in grey indicate significant differences in performance between groups **c-e)** Post assessment performance (post-R vs post-NR) comparing Rew and NoRew groups on **c)** Day1, **d)** Day2 and **e)** Day7. Shaded regions/error bars represent SEM.

Importantly, the result also suggests that the reward groups may further improve FI performance over the course of training which aligns with previous results showing that learning to transition between sequential reaching movements faster is a reward sensitive training-depending process^10^.

Assessing performance across post assessment, we conducted separate mixed-effect ANOVAs for each day and found for each day a main effect for reward only on Day2 (Day1: F = 3.31, p = 0.0724, η^2^ = 0.77; Day2: F = 5.10, p = 0.0264, η^2^ = 0.84; Day7: F = 2.06, p = 0.1543, η^2^ = 0.67). Yet, we found timepoint to be significant on all days (Day1: F = 23.34, p < 0.0001, η^2^ = 0.96; Day2: F = 27.60, p < 0.0001, η^2^ = 0.97; Day7: F = 65.33, p < 0.0001, η^2^ = 0.99). Similalry to vel_min_, we found a significant reward x timepoint interaction only for Day3 (Day1: F = 0.28, p = 0.5995, η^2^ = 0.22, Figure 5c; Day2: F = 0.49, p = 0.4856, η^2^ = 0.33, Figure 5d; Day7: F = 13.72, p < 0.0001, η^2^ = 0.93, Figure 5e). Post-hoc analysis revealed that FI scores changed significantly from post-R to post-NR only for the NoRew groups (Z = 2.54, p = 0.0322, η^2^ = 0.27, Figure 5e) but not for the Rew groups (Z = 1.00, p = 0.3164, η^2^ = 0.10). Additionally, performance was significantly different when comparing the Rew and NoRew groups during post-NR (Z = 2.41, p = 0.0322, η^2^ = 0.25) but not post-R (Z = 1.02, p = 0.3164, η^2^ = 0.11). Similarly to vel_min_, these results suggest that FI performance was less reward dependent and became more stable in the Rew groups on Day7.

## Discussion

Previous work was able to demonstrate motor sequence learning is driven by distinct behavioural invigoration processes^10^. Specifically, it was shown that a monetary incentive led to an increase in motor vigour through increases in vel_peak._ In contrast, performance-based feedback enhanced movement fusion indexed as increases in vel_min_ and FI. In the present study, we sought to investigate whether these behaviourally distinct reward-based processes rely on dissociable dopaminergic mechanisms. Our results showed that the D2 antagonist haloperidol modulated motor sequence learning as seen in drug-specific changes in MTs over the course of Training on Day1. Specifically, it appeared that while both Halo groups were significantly faster during early Training, we detected an increase in MTs over the course of Training. This finding converges with previous research showing that haloperidol lowers the willingness to exert effort^48–50^ leading to a decrease in motor sequence learning (i.e., MTs). Importantly, we found that this slowing in MTs was driven by a selective impairment of the reward-based effects on motor vigour as seen in reduced vel_peak_ in the Halo-Rew group. In contrast, we found that haloperidol had no detrimental effect on reward-based increases in movement fusion as measured in changes in vel_min_ and FI. Therefore, our results highlight that the motivational (driven by monetary incentives) and informational (driven by performance-based feedback) properties of reward are differentially affected by a DA manipulation suggesting dissociable underlying mechanisms.

### Haloperidol selectively affects the reward-based effects on motor vigour

Our results showed that the D2 antagonist haloperidol impaired the reward-based effects on vel_peak_ during a sequential reaching task. Specifically, we found that vel_peak_ were lower in the Halo-Rew group compared to Placebo-Rew group, while no differences between the NoRew were found. These results suggest that haloperidol reduced the reward-based effects on motor vigour which aligns with previous work showing that DA plays a role in the integration of efforts and rewards^48^. Specifically, research demonstrated that a DA blockage reduced the willingness to choose effortful options (i.e., increase motor vigour) to maximise rewards^48,50–53^. These results complement recent behavioural work showing that reward increases vel_peak_ in a simple reaching task. Crucially, decreases in MTs via increases in vel_peak_ were transient in nature and disappeared once reward was removed^13,14^. This has been interpreted as reward (i.e., a monetary incentive) paying the energetic cost of increased motor vigour. Consequently, once reward was removed there was no incentive to maintain increased motor vigour^12,18^. Here we replicate findings demonstrating that a DA blockage leads to a similar behavioural result (i.e., decrease in motor vigour) by altering the effort-reward trade-off. Interestingly, we found that haloperidol did not affect vel_peak_ in the Halo-NoRew when compared to Placebo-NoRew. This is somewhat surprising considering Parkinson’s disease (P) patients OFF medication show a reduced willingness to exert effort independent of the reward context^52,53^. It is currently unclear what could explain this dichotomy however the relative specific effect of haloperidol on D2-receptors is a possibility.

### Haloperidol does not impair movement fusion

In the present task, MTs could be reduced via two independent strategies: 1) an increase in vel_peak_ through and increase in motor vigour and 2) an increase in fusion measured as an increase in vel_min_ and/or alternatively as an increase in FI. By reducing dwell times of reaching movement transitions, movement fusion allows for a faster but also smoother execution making the overall movement more efficient^10,54^. Therefore, movement fusion represents a hallmark of motor sequence learning and has been shown to be often impaired in clinical populations such as stroke and PD patients^1–3^. Based on evidence from clinical studies investigating this aspect of motor sequence learning (i.e., fusion, chunking, coarticulation), DA has been implicated to underlie this process^1,45,46^. Converging results come from animal work (i.e., rodents^40,41,43^ and monkeys^42^) and studies conducted with healthy humans^44^. However, here we did not find that a DA manipulation using the D2 antagonist haloperidol affected motor sequence learning. Specifically, we did not observe that haloperidol impaired the reward-based effects on vel_min_ and FI. Instead, we found that both Rew groups exhibited reward-based increases in vel_min_ and FI leading to more pronounced movement fusion when compared to the NoRew groups. The lack of a haloperidol effect on fusion in either Halo group further indicates that fusion may not rely strongly on the availability of D2 DA. This finding diverges from previous work in monkeys showing that raclopride, another D2 antagonist with high D2 binding, impaired chunking^42,55^. However, it is unclear if raclopride and haloperidol lead to a similar D2 blockage and whether a prolonged use of either of them over multiple training days with drug intake increases the drug-specific effects on fusion.

In summary, our results highlight that in humans the reward-based effects on motor vigour and movement fusion are differentially affected by dopaminergic manipulation suggesting dissociable underlying mechanisms.

## Methods

### Participants

95 participants (42 males; age range 18 - 42) were recruited to participate in three experiments, which had been approved by the local research ethics committee of the University of Birmingham. All participants were novices to the task paradigm and were free of motor, visual and cognitive impairment. Potential participants were pre-screened and were only invited to the medical exam if they met the following criteria: 1) naïve to the task paradigm ;2) 18–45 years old; 3) no self-reported history of medical disorders; 4) normal or corrected-to-normal vision; 5) no drug allergies; 6) currently taking no medication that interfere with the absorption of haloperidol. Suitable participants were then evaluated by a medical doctor, who reviewed their medical history, evaluated an electrocardiogram taken at rest and took a blood pressure reading. Participants who received medical approval were then scheduled for all experimental sessions. Most participants were self-reportedly right-handed (N = 7 left-handed participants) and gave written informed consent prior to the start of the experiment. Participants were remunerated with money (£18/hour) and were able to earn additional money during the task depending on their performance. Before the start of the experiment, participants were pseudo-randomly allocated to one of the available groups.

### Experimental Protocol, Randomisation and Blinding Protocol

In this study, we sought to investigate whether DA modulates the reward-based improvement of movement vigour and/or movement fusion. To this end, participants were randomly allocated to one of four groups: haloperidol with reward-based feedback (Halo-Rew, N = 25), haloperidol without reward (Halo-NoRew, N = 24), placebo with reward-based feedback (Ctrl-Rew, N = 23) and placebo without reward (Ctrl-NoRew, N = 23) after the medical assessment (Figure 1c). Due to the required testing environment this study was single-blind, and both the medical doctor and examiner were aware of the drug group allocation (haloperidol vs placebo). However, to reduce bias all participants were told that they would receive either a placebo tablet or the active drug (haloperidol). Similarly, all participants had to complete a health check on the day of drug/placebo intake (Day1) and were checked by the medical doctor in intervals of 1h throughout Day1. Additionally, all task instructions were displayed on screen instead of communicated verbally to further reduce bias. The administration of both the active drug and the placebo tablet was performed by the medical doctor.

On the day of drug/placebo intake (Day1), participants either received 2.5mg of haloperidol (2 × 0.5mg and 1 × 1.5mg tablet) or three lactose tablets of the same white colouring. In each case, participants were handed an envelope containing either the active drug or placebo and were asked to close their eyes during intake. Haloperidol is a D2-receptor antagonist that shows a limited affinity to D1 receptors and has superior in vivo D2 binding. In addition, it blocks DA D2 binding in the basal ganglia (BG) but not in the prefrontal cortex and as such can be considered to selectively modulate DA levels within the BG pathway^56^. To coincide with the peak plasma concentration, participants were asked to wait in the lab for 120min before engaging in the motor task^56^. The chosen dose of 2.5mg of haloperidol and the waiting time of 2h were similar to previous studies that were able to observe drug-related behavioural and neurophysiological effects of haloperidol^56,57^. After the waiting period, participants were asked to complete the motor task (see Task Design and Task Protocol for details) and upon completion were yet again checked by the medical doctor. On Day1 participants were additionally asked to complete a questionnaire to report their perceived levels of fatigue and attention. We added this self-report because while haloperidol is a known D2 antagonist it may also act as sedative which could influence results. 42 participants (Halo = 21; Placebo = 21) completed the questionnaire (which was added while the trial was already ongoing) and answered the following questions: ‘Please rate yourself with regards to: 1) Attention and 2) Fatigue’. The scale ranged from 1-8 with lower numbers indicating lower levels of attention and fatigue. Participants were scheduled to return to the lab 24h later (Day2) to complete the same motor task again; this time without any drug/placebo manipulation. However, participants received the same feedback as during Day1 (i.e., with or without reward-based feedback during Training). Lastly, participants engaged in a short version of the motor task one week after the initial session (Day7) and were subsequently debriefed. **Experimental apparatus:** The experiment was performed using a Polhemus 3SPACE Fastrak tracking device (Colchester, Vermont U.S.A; with a sampling rate of 110Hz). Participants were seated in front of the experimental apparatus which included a table, a horizontally placed mirror 25cm above the table and a screen (Figure 1a). A low-latency Apple Cinema screen was placed 25cm above the mirror and displayed the workspace and participants’ hand position (represented by a green cursor – diameter 1cm). On the table, participants were asked to perform 2-D reaching movements. Looking into the mirror, they were able to see the representation of their hand position reflected from the screen above. This setup effectively blocked their hand from sight. The experiment was run using MATLAB (The Mathworks, Natwick, MA), with Psychophysics Toolbox 3.

### Task design

Participants were asked to hit a series of targets displayed on the screen (Figure 1a, b). Four circular (cm diameter) targets were arranged around a centre target (‘ ia target’). Starting in the via target, participants had to perform eight continuous reaching movements to complete a trial. Target 1 and 4 were displaced by 10cm on the *y-*axis, whereas Target 2 and 3 were 5cm away from the via target with an angle of 126 degrees between them (Figure 1b). To start each trial, participants had to pass their cursor though the preparation box (2×2cm) on the left side of the workspace, which triggered the appearance of the start box (2×2cm) in the centre of the screen. After moving the cursor into the start box, participants had to wait for 1.5s for the targets to appear. This ensured that participants were stationary before reaching for the first target. Target appearance served as the go-signal and the start box turned into the via target (circle). Upon reaching the last target (via target), all targets disappeared, and participants had to wait for 1.5s before being allowed to exit the start box to reach for the preparation box to initiate a new trial. Participants had to repeat a trial if they missed a target or performed the reaching order incorrectly. Similarly, exiting the start box too early either at the beginning or at the end of each trial resulted in a missed trial.

### Reward structure and feedback

Participants in the Rew groups were informed that faster MTs would earn them more money. Reward trials were cued using a visual stimulus prior to the start of the trial (Figure 1d left). Once participants moved into the preparation box, the start box appeared in yellow (visual stimulus). In contrast, participants that were in a NoRew groups were told to move as fast and accurately as possible and here the start box remained black. Performance feedback was provided after completing a trial while participants moved from the start box to the preparation box to initiate a new trial. Feedback was displayed on the top of the screen (i.e., ‘ p out of 5p’). We used a closed-loop design to calculate the feedback in each trial. To calculate this, we included the MT values of the last 20 trials and organised them from fastest to slowest to determine the rank of the current trial within the given array. A rank in the top three (<= 90%) returned a value of 5p, ranks >= 80% and <90% were valued at 4p; ranks >=60% and <80% were awarded 3p; ranks >=40% and < 60% earned 2p while 1p was awarded for ranks >=20% and < 40%. A rank in the bottom three (<20%) returned a value of 0p. When participants started a new experimental block, performance in the first trial was compared to the last 20 trials of the previously completed block.

### Task Protocol

The main experiment included four experimental parts: Baseline, Training, a post assessment with reward and one without (Figure 1f). Participants completed this protocol on Day1 and Day2, while only completing the two post assessments on Day7. Additionally, a learning block was scheduled prior to the start of the main experiment on Day1. Furthermore, a secondary task was included in this task design, which asked participants to press a force sensor with the index finger of their non-dominant hand. Participants were told to apply pressure in response to an audio signal that changed in amplitude, with higher amplitudes requiring increased force and vice versa. However, the analysis of these secondary-task trials which were scheduled on every 10^th^ trial during training and every 5^th^ trial during the remaining parts are excluded from the analysis presented here.

#### Learning

We included a learning phase prior to the start of the experiment on Day1 so that participants could memorise the reaching sequence (Figure 1b). After completing 20 learning trials, participants moved on to the main experiment.

#### Baseline

Participants in both groups completed 10 baseline trials, which were used to assess whether there were any pre-training differences between groups. oth groups were instructed to ‘mo e as fast and accurately as possible’, while no performance-based feedback was given at the end of each trial. *Training:* Participants in the Rew groups were informed that during this part they would be able to earn money depending on how fast they complete each trial (200 reward trials). In contrast, participants in the no reward group engaged in 200 no reward trials and were again instructed to move as fast and as accurately as possible.

#### Post assessments

Participants from both groups were asked to complete two post assessments (20 trials each on Day1 and Day2 and 25 trials on Day7); one with reward trials (post-R) and one with no reward trials (post-NR). The order was counter-balanced across participants.

Consequently, participants completed three experimental sessions (Day1, Day2 and Day7) as well as a medical assessment prior to the start of the study (Figure 1f).

### Dual Task

Additionally, a secondary task was included in this task design, which asked participants to press a force sensor with the index finger of their non-dominant hand. Participants were told to apply pressure in response to an audio signal that changed in amplitude, with higher amplitudes requiring increased force and vice versa. However, the analysis of these secondary-task trials which were scheduled on every 10^th^ trial during training and every 5^th^ trial during the remaining parts are excluded from the analysis presented here.

### Data analysis

Analysis code is available on the Open Science Framework website, alongside the experimental datasets at: https://osf.io/62wcz/. The analyses were performed in Matlab (Mathworks, Natick, MA) and JASP.

### Movement time (MT)

MT was measured as the time between exiting the start box and reaching the last target. This excludes reaction time, which describes the time between target appearance and when the participants’ start position exceeded 2cm.

### Maximum and minimum velocity

Through the derivative of positional data (*x, y*), we obtained velocity profiles for each trial which were smoothed using a gaussian smoothing kernel (σ =). he velocity profile was then divided into segments representing movements to each individual target (8 segments) by identifying when the positional data was within 2cm of a target. We measured the maximum velocity (*v*_*max*_) of each segment by finding the maximum velocity:

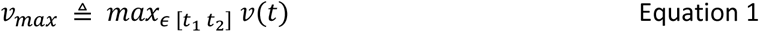

Where *v(t)* is the velocity of segment *t*, and *t*_*1*_ *and t*_*2*_ represent the start and end of segment *t* respectively. Similarly, minimum velocities (*v*_*min*_) were determined by measuring the minimum velocity when participants were inside a target (7 targets) using:

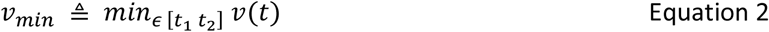

The individual maximum and minimum velocities were then averaged for each trial.

### Fusion index (FI)

Fusion describes the blending together of individual motor elements into a singular smooth action. This is represented in the velocity profile by the stop period between the two movements gradually disappearing and being replaced by a single velocity peak (Figure 4a, b) ^19–21^. To measure fusion, we compared the mean maximum velocities of two sequential reaches with the minimum velocity around the via point. The smaller the difference between these values, the greater coarticulation had occurred between the two movements (Figure 4b) ^58^. We calculated movement fusion by:

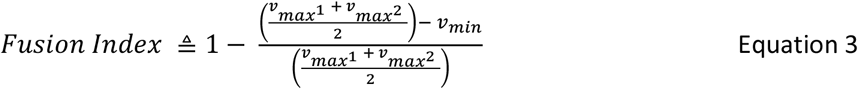

with *v*_*max*_^*1*^ and *v*_*max*_^*2*^ representing the velocity peak value of two reaching movements, respectively, and *v*_*min*_ representing the minimum value between these two points. We normalised the obtained difference, ranging from 0 to 1, with 1 indicating a fully fused movement. Given that in this task seven transitions had to be completed, the maximum FI value was 7 in each trial.

### Statistical analysis

Assessing Baseline performance, we found that 3 participants exhibited >8.5s to complete a trial. We decided to remove these participants from further analysis after conducting an outlier analysis and comparing their mean (10.03s ± 1.0) to the rest of the participants (5.37s ± 1.1). In total, 2 participants from the Halo-NR group and 1 further participant from the Placebo-NR group were removed. This changed the group sizes with Halo-R (N=25), Halo-NR (N=21), Placebo-R (N=24) and Placebo-NR (N=22). Furthermore, trials that took longer than >10s were excluded from further analysis (N = 11). Finally, all dual task trials were removed from the analysis. Dual task trials were scheduled on every 10^th^ trial during training and every 5^th^ trial during the remaining parts which meant that for each participant 2 trials were removed from Baseline, 20 from Training and 4 and 5 during the post assessments on Day1/2 and Day7 respectively.

Using one-sample Kolmogorov–Smirnov tests to test our data for normality we found that performance during Baseline was nonparametric. We, therefore, calculated the median of participants’ performance and used a Kruskal Wallis test to determine performance differences at Baseline. Furthermore, we used linear mixed-models (LMM) to assess statistical significance of our Training results. We carried out separate analyses for Training on Day1 (Training1) and Day2 (Training2). LMMs were conducted in JASP and included three fixed effects: 1. reward (1 = Reward; 2 = No Reward), 2. drug (1 = haloperidol; 2 = placebo) and trial number (1:180). By default, all the fixed effects and their interactions are included in the model. Therefore, the model included all possible interaction terms between the three main effects (i.e., *Reward *Drug, Reward *Trial Number, Drug *Trial Number, Reward *Drug *Trial Number*). Individual differences were accounted for by including Participant (1:92) as a random effect. By default, all the random effects corresponding to the fixed effects are included which corresponds to the “maximal random effects structure justified by the design"^59^. Therefore, the model included random slopes for all fixed and random effects combinations (i.e., *Reward *Drug, Reward *Trial Number, Drug *Trial Number, Reward *Drug *Trial Number*). However, all random slopes involving Reward and Drug were removed from analysis because they do not vary within a participant (i.e., they refer to a group). In contrast, Trial was included as a random slope which improved the model as seen in a lower BIC. Removing Trial as a random slope increased the BIC and we therefore used the following model for all LMMs presented in the paper:

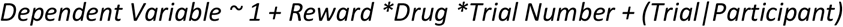

Additionally, the model terms were tested with the Satterthwaite method and a restricted maximum likelihood method was used to fit the model. How to best investigate significant interactions observed within LMM has been a heated topic^60^. Here, we used a row-shuffle non-parametric permutation test to assess whether groups differ across training. To this end, we computed 1000 permutations for each trial during which participants were row-shuffled into groups. We obtained the t-statistic of each permutation and subsequently determined whether the real t-statistic is statistically different from the permuted ones (p < 0.05). This allowed us to investigate during which trials groups were exhibiting significantly different performance. Additionally, mixed-model ANOVAs were used to assess statistical significance during the post assessments, with reward (reward vs no reward), drug (haloperidol vs placebo) as between factors and timepoint (post-R vs post-NR) as a within factor. We used one-sample Kolmogorov–Smirnov tests to test our data for normality and found that all measures were nonparametric. Median values were therefore used as input in all mixed-model ANOVAs (similar to^10,61^). Wilcoxon tests were employed when a significant interaction and/or main effects were reported. The results were corrected for multiple comparisons with false discovery rate [fdr_bh(stats, ‘alpha’,. 5) in MA LA ^62^]. Therefore, the P values presented in results have been adjusted to the number of comparisons conducted (comparing each group with all others, which amounts to N = 4).

## Acknowledgements

This work was supported by the European Research Council starting grant: MotMotLearn (637488)

## Supplement

**Supplementary Table 1:** Model Summary.

| Model | R | R<sup>2</sup> | Adjusted R<sup>2</sup> | RMSE | R<sup>2</sup> Change | F Change | df1 | df2 | p |
| --- | --- | --- | --- | --- | --- | --- | --- | --- | --- |
| 1 | 0.000 | 0.000 | 0.000 | 1.157 | 0.000 |  | 0 | 33113 |  |
| 2 | 0.804 | 0.647 | 0.647 | 0.688 | 0.647 | 60561.856 | 1 | 33112 | < .001 |
| 3 | 0.859 | 0.737 | 0.737 | 0.593 | 0.091 | 11458.614 | 1 | 33111 | < .001 |
| 4 | 0.920 | 0.846 | 0.846 | 0.455 | 0.108 | 23252.379 | 1 | 33110 | < .001 |

**Supplementary Table 2:** Coefficients.

| Model |  | Unstandardized | Standard Error | Standardized | t | p | Collinearity Statistics |  |
| --- | --- | --- | --- | --- | --- | --- | --- | --- |
|  |  |  |  |  |  |  | Tolerance | VIF |
| 1 | Intercept | 3.822 | 0.006 |  | 601.085 | < .001 |  |  |
| 2 | Intercept | 7.250 | 0.014 |  | 502.346 | < .001 |  |  |
|  | Peak | -0.110 | 4.478e -4 | -0.804 | -246.093 | < .001 | 1.000 | 1.000 |

**Supplementary Table 2: Coefficients**
| Model |  | Unstandardized | Standard Error | Standardized | t | p | Collinearity Statistics |  |
| --- | --- | --- | --- | --- | --- | --- | --- | --- |
|  |  |  |  |  |  |  | Tolerance | VIF |
| 3 | Intercept | 6.591 | 0.014 |  | 474.868 | < .001 |  |  |
|  | Peak | -0.071 | 5.346e -4 | -0.515 | -132.079 | < .001 | 0.521 | 1.918 |
|  | Min | -0.081 | 7.600e -4 | -0.418 | -107.045 | < .001 | 0.521 | 1.918 |
| 4 | Intercept | 8.744 | 0.018 |  | 494.570 | < .001 |  |  |
|  | Peak | -0.134 | 5.823e -4 | -0.976 | -229.602 | < .001 | 0.258 | 3.875 |
|  | Min | 0.189 | 0.002 | 0.969 | 101.219 | < .001 | 0.051 | 19.652 |
|  | CI | -1.379 | 0.009 | -1.166 | -152.487 | < .001 | 0.080 | 12.539 |

